# Modelling antibody structures at the speed of language

**DOI:** 10.64898/2026.06.03.729879

**Authors:** Isaac Ellmen, David Errington, Matthew I.J. Raybould, Charlotte M. Deane

## Abstract

Protein structure prediction is currently substantially slower than obtaining sequence representations of proteins. This leads to most property prediction methods relying solely on trivial or learned sequence embeddings. However, contemporary structure prediction and sequence models are both based on Transformers, and structure prediction models often have fewer parameters, suggesting that there might be domains where accurate structure prediction adds no practical overhead to sequence-only modelling. Here, we demonstrate this can be achieved for adaptive immune proteins by introducing FlashABB, which predicts highly accurate antibody structures, and does so faster than even modestly-sized language models can embed sequences. As a component of FlashABB, we develop Flashpoint Attention, a fast and linear memory analog of Invariant Point Attention. To our knowledge, FlashABB is the first example of a model that accurately predicts protein structure faster than protein language models can generate embeddings, enabling efficient access to 3D information without the need for precomputed structures. Using FlashABB, we develop methods for predicting antibody stability and developability which can be scaled to repertoires of millions of sequences. Our results show how the computational bottleneck of protein structure prediction can be removed in some real-world cases.

The code and model weights for FlashABB are available on GitHub: https://github.com/oxpig/FlashABB

## 1 Introduction

Machine learning-based structure prediction has enabled substantial advances in protein design [1, 2] and property prediction [3, 4], including in applications like therapeutic antibody design [5–8]. However, protein structure prediction is computationally intensive, which limits its use on large-scale sequence datasets such as antibody repertoires. Models like AlphaFold2 (AF2) usually take minutes or hours [9, 10], whereas protein language models (PLMs) like ESM2 typically generate protein embeddings in a fraction of a second. In some ways this is unintuitive, as at their core, structure models such as AlphaFold2 and sequence models such as ESM2 are both Transformers and use very similar architectures. In fact, structure models are typically much smaller than sequence models. The smallest ESM2 model has a hidden dimension of 320 whereas the AF2 structure module has a hidden dimension of 128 [9, 10].

The reasons for the difference in efficiency are complex, but largely stem from three expensive modifications:

1. The analysis of coevolutionary information and subsequent processing of the pair representation, which typically adds **minutes**.
2. Molecular dynamics refinement of generated structures, which adds on the order of **seconds**.
3. Inefficient kernels for Invariant Point Attention compared to standard scaled dot-product attention, which adds on the order of **10s of milliseconds**.

In this work, we close the efficiency gap between structure and sequence modelling for antibodies, an important class of proteins. Our new method, FlashABB can predict about 200 structures per second on an NVIDIA RTX A2000. This is about 1000x as fast as ABodyBuilder2, about 3x as fast as loading a PDB structure with BioPython, and about 2x as fast as AbLang2, an antibody language model based on ESM2-35M [6, 11, 12]. The efficiency of FlashABB enables the inclusion of structure with negligible overhead for antibody sequence models.

Most Transformers perform fully-connected attention, which means that they must perform *n*^2^ inner products and generate an *n* × *n* attention matrix [13]. However, modern attention implementations such as FlashAttention leverage the fact that it is faster for GPUs to compute inner products than to load values from memory, which permits very fast attention implementations which only use linear memory [14, 15]. Similarly, Rabe and Staats [16] proposed an attention method which uses gradient checkpointing to compute self-attention in 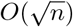 memory.

Fast, memory-efficient methods are used commonly in the pair processing of methods like OpenFold and Boltz [17, 18], however these methods still require quadratic memory and do not fully leverage inner product attention in Invariant Point Attention (IPA). Concurrently with our work, Liu et al. [19] proposed a similar efficient IPA analog to ours. For a complete comparison of the methods, see Appendix A.

In this work we demonstrate our method’s ability to rapidly model protein structures by focusing on antibodies. The reason for this is three-fold. First, antibodies are important components of the immune system and antibody therapies are ubiquitous in fields such as oncology [20]. Second, antibody variable regions are notoriously difficult to model. For example, AlphaFold3 only achieves an RMSD of 2-3Å on the CDRH3 region of antibodies, which is substantially worse than experimental accuracy [21, 22]. Third, antibodies develop independently in each person which means that there are not useful evolutionary clues in sequencing data [5, 6, 23]. This enables ABodyBuilder(2/3) (ABB(2/3)) to achieve state of the art accuracy in modelling antibody loops using a trivial pair representation [6, 24]. In this work we demonstrate that the trivial pair representation can be removed and the features can be generated “on the fly”, enabling a far more efficient attention mechanism.

## 2 Results

In this section, we demonstrate that our Flashpoint Attention algorithm provides a significantly more efficient alternative to standard IPA under a simple pair representation. We use it to create FlashABB, a super-fast antibody structure prediction method and show that it produces similar backbone structures to ABodyBuilder2/3. Finally, we use FlashABB to predict (1) stability and (2) therapeutic developability properties. All structures and properties can be predicted in milliseconds, which enables repertoire-scale antibody structural analysis.

### 2.1 Flashpoint Attention performance

We tested the speed and memory requirements for our Flashpoint Attention (FPA) implementation compared to the OpenFold IPA implementation [17]. We ran each attention method 100 times for a randomly chosen antibody (sequence listed in Appendix B) replicated over a batch size of 32, and report the median time and memory used. The results are shown in Figure 2. In both the forward and backward pass, our method was over 6x as fast and used over 10x less memory.

**Fig. 1.**
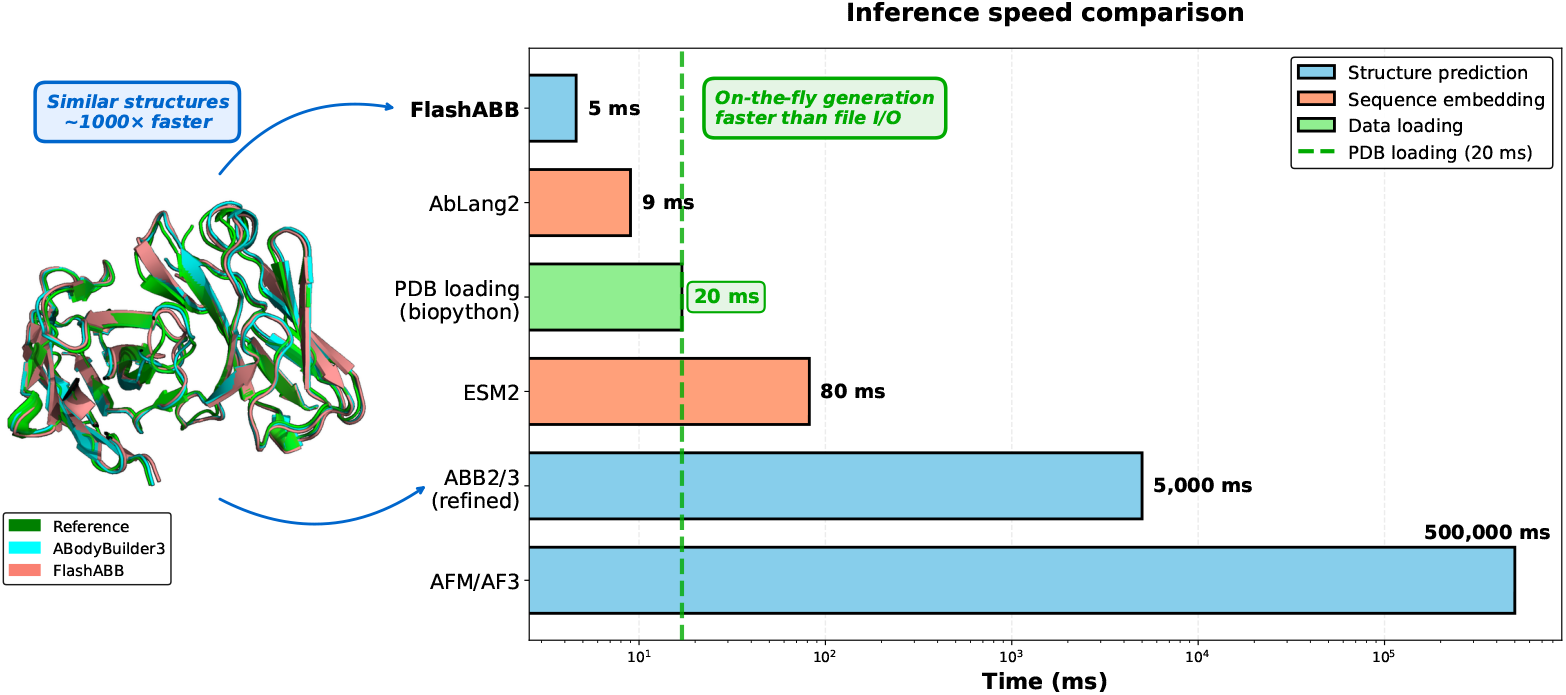
Inference time per antibody for various models. ESM2 is the 650M parameter model, PDB loading is the average time to load and parse an antibody structure with BioPython, ABB2/3 are ABodyBuilder models, and AFM/AF3 are AlphaFold-Multimer and AlphaFold3. Times are approximate, but FlashABB is about twice as fast as AbLang2, an antibody language model and over 10,000 times as fast as AlphaFold models with MSA construction.

**Fig. 2.**
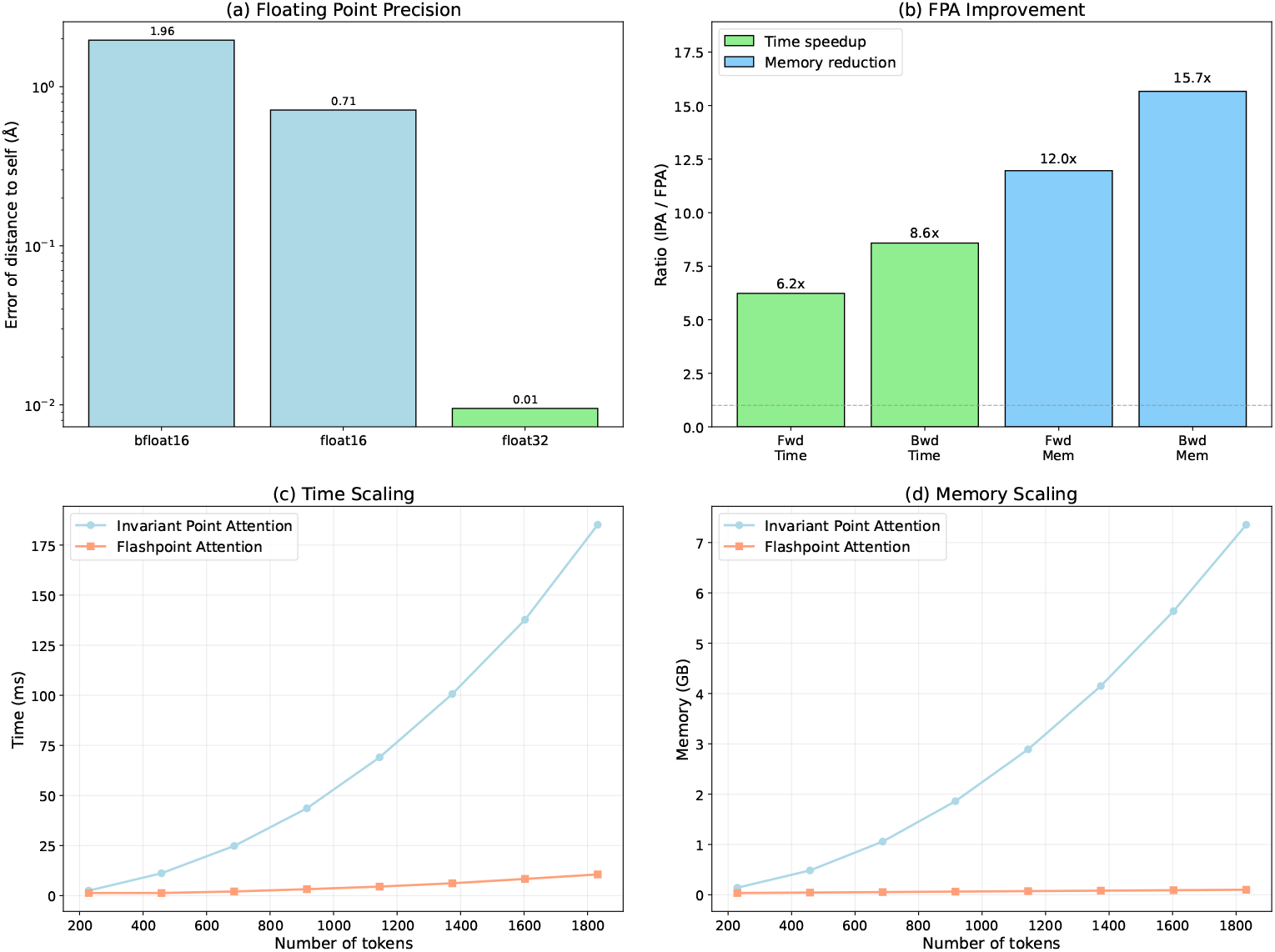
Properties of FPA. (a) Distance computation accuracy under different levels of precision. 32 bit floats can compute distance to within 0.01Å whereas 16 bit floats meaningfully suffer from catastrophic cancellation. (b) Average time and memory for a batch of 32 antibodies run through FPA and IPA. FPA is substantially faster and more memory efficient. Time (c) and memory (d) as a function of sequence length for FPA and IPA. FPA still uses quadratic time but only linear memory.

To verify the scaling properties of our FPA implementation we tested on longer sequences. The results are shown in Figure 2. As expected, the time per sequence remains substantially lower over longer sequences but scales quadratically, while the memory scales linearly. This is particularly notable for very long sequences, where IPA would be intractable even with many GPUs, whereas FPA’s memory footprint remains negligible (*∼*100MB).

### 2.2 FlashABB accuracy

We tested our FPA implementation by training on antibody structure prediction using the sets described in Methods Section 4.4. Results are shown in Table 1 and an example structure is shown in Figure 3. ABB(2/3) each use a molecular dynamics refinement step which improves RMSD values across all regions [24]. To compare the machine learning model performance, we reproduce the results for ABB3 without the structural refinement. Our model is broadly comparable to ABB3 in all regions across the antibody backbone. We observed that our training loss was slightly worse due to a lower Frame Aligned Point Error (FAPE) term, however this had no impact on the backbone accuracy. Benchmarking the performance of frontier models like Boltz, Chai, and AlphaFold3 is difficult due to the tunability of hyperparameters like MSA construction and diffusion steps as well as uncertainty in data leakage from training, however most studies have found that these typically achieve CDRH3 accuracies of 2-3Å [18, 21, 22, 25, 26].

**Table 1.** Mean backbone RMSD (Å) by CDR and framework region.

|  | CDRH1 | CDRH2 | CDRH3 | Fw-H | CDRL1 | CDRL2 | CDRL3 | Fw-L |
| --- | --- | --- | --- | --- | --- | --- | --- | --- |
| ABB2* | 0.84 | 0.73 | 2.54 | 0.56 | 0.55 | 0.36 | 0.88 | 0.53 |
| ABB3* | 0.87 | 0.70 | 2.42 | 0.58 | 0.61 | 0.39 | 0.93 | 0.58 |
| ABB3 (unrefined) | 0.78 | 1.01 | 2.62 | 0.65 | 0.76 | 0.49 | 0.97 | 0.63 |
| FlashABB | 0.80 | 1.05 | 2.59 | 0.67 | 0.71 | 0.43 | 0.98 | 0.64 |

\*As reported in Kenlay et al. [24]

**Fig. 3.**
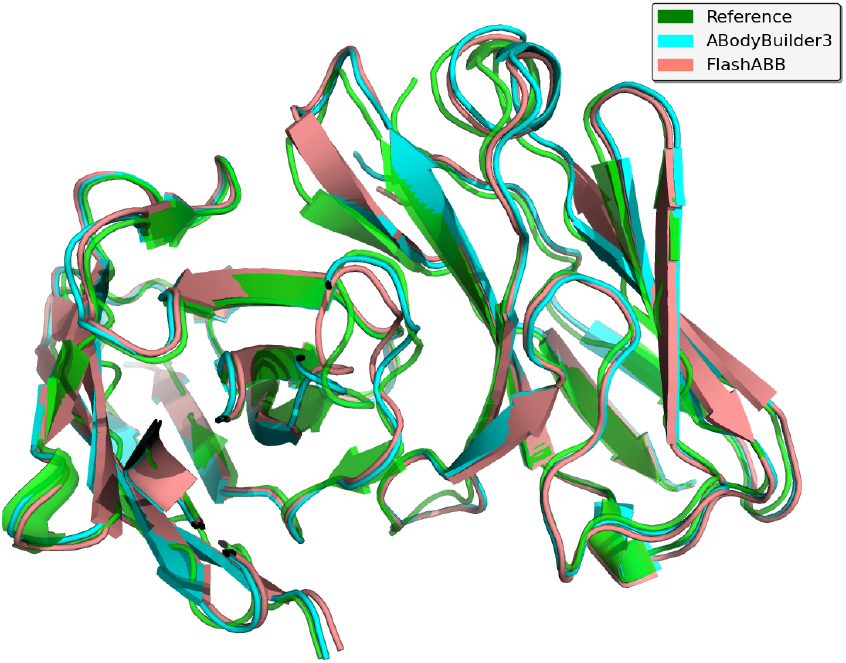
Ribbon representation of a randomly selected antibody structure (PDB: 8DG9). Green is the ground truth structure, cyan is the refined ABodyBuilder3 prediction, and salmon is our unrefined prediction. The unrefined prediction may contain minor stereochemical errors but adopts a very similar global fold.

### 2.3 Stability enrichment at scale

Inverse folding likelihoods are known to correlate well with protein stability [3]. Classically, the compute bottleneck for using inverse folding models for stability prediction is protein structure prediction. We tested whether FlashABB-predicted structures could be used to condition ProteinMPNN for super-fast antibody stability prediction. We ran FlashABB and ProteinMPNN on all antibodies in the set from Widatalla et al. [27] which is comprised of human antibodies from Shehata et al. [28] and therapeutic antibodies from Jain et al. [29]. As a control, we computed likelihoods using pIgGen, an autoregressive paired antibody language model [30]. We found that the FlashABB+ProteinMPNN-derived likelihoods correlated well with stability in both therapeutic and natural antibodies (Figure 4), which indicates that it has learned sound biophysical principles which determine stability. Conversely, pIgGen likelihoods correlated well with natural antibodies but had almost no correlation with therapeutic antibodies. This is probably because germline antibodies tend to be more stable and are better predicted by language models [11, 28].

**Fig. 4.**
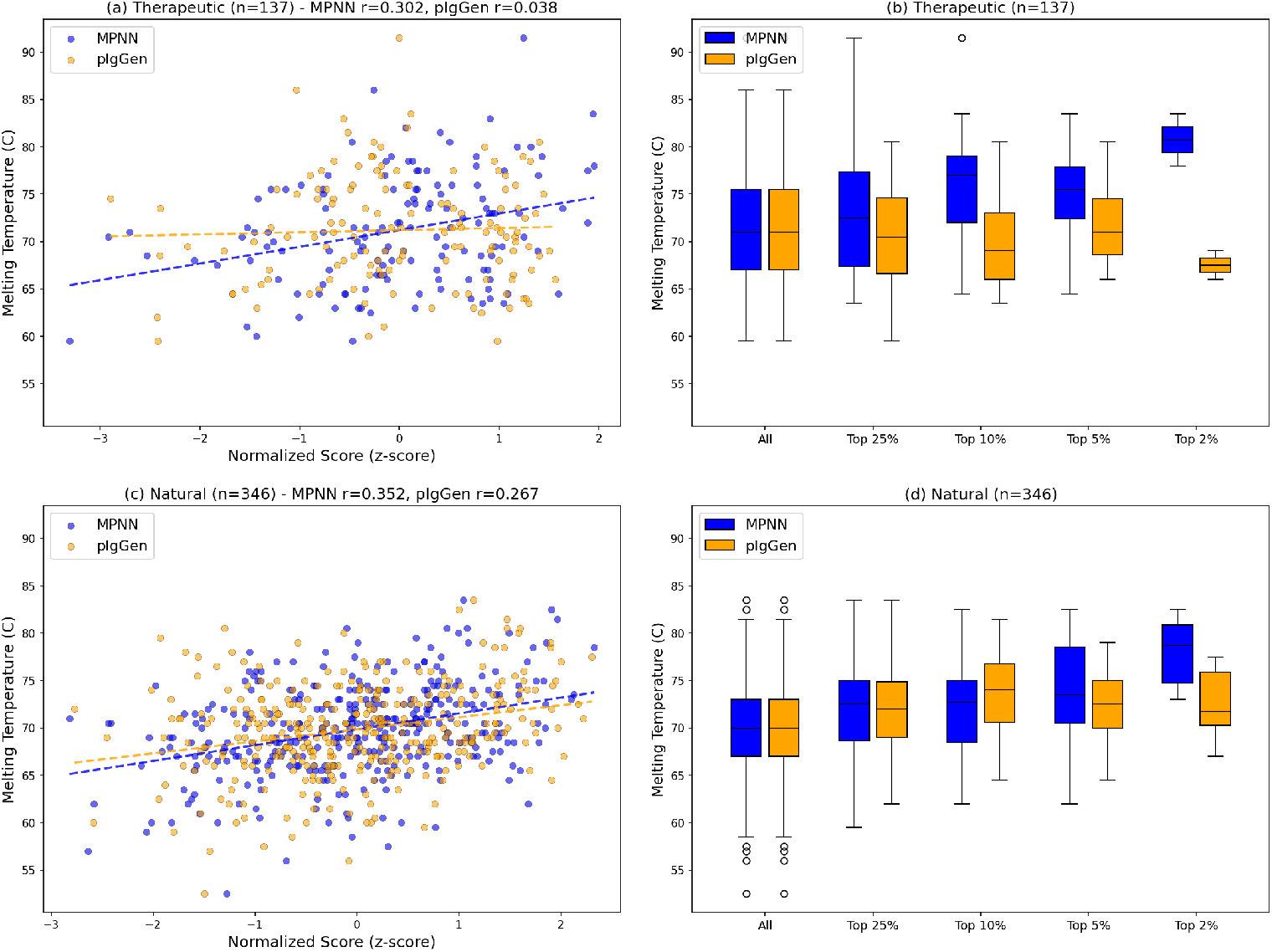
Log likelihood vs melting temperature for FlashABB+ProteinMPNN (labelled MPNN) and pIgGen. Both methods have a positive correlation over natural antibodies because of the language model preference for germlines. FlashABB+ProteinMPNN retains a good correlation with stability over therapeutic antibodies which requires an internal representation of the physicochemistry. This enables stability enrichment. The model can estimate stability for *∼*50 sequences per second and the median Tm values of the therapeutic MPNN-enriched subsets are 71, 72.5, 77, 75.5, and 81 °C.

We next investigated whether FlashABB+ProteinMPNN could be used for stability enrichment. We looked at the median melting temperatures of the top 25%, 10%, 5%, and 2% of antibodies in our test set as scored by FlashABB+ProteinMPNN. The top 10% of antibodies had a median stability improvement of 6°C. Additionally, because FlashABB removes the structure prediction bottleneck from the approach, we were able to estimate stability for *∼*50 antibodies per second on a single NVIDIA RTX A2000 GPU. This means this approach could practically be used to enrich for stability in libraries of hundreds of millions of antibodies.

### 2.4 FlashTAP

To demonstrate the utility of our FlashABB-SSS seq2seq model, we finetuned it for property prediction of TAP developability properties. The Therapeutic Antibody Profiler (TAP) is a tool for predicting developability issues in therapeutic antibody candidates [7, 31]. Four of the TAP properties are computed on antibody structures which can make it expensive to run at the scale of repertoires. We trained three models, a baseline onehot encoding model, a finetuned language model based on AbLang2, and a finetuned version of FlashABB-SSS. We evaluated each model’s mean average error (MAE) for each property. TAP properties are also used to assign ‘flags’ for likely developabilty issues based on predefined thresholds. We computed the precision-recall curves for predicting flags while setting different thresholds for each model to evaluate their general-purpose utility at identifying issues. The MAE and area under the precision-recall curve (AUPRC) values are plotted in Figure 5. While AbLang2 provided a strong baseline, FlashABB-SSS was the best underlying model in call cases. The difference was particularly pronounced for SFvCSP AUPRC on the therapeutic set, which may be due to a better generalized prediction based on true structural understanding.

**Fig. 5.**
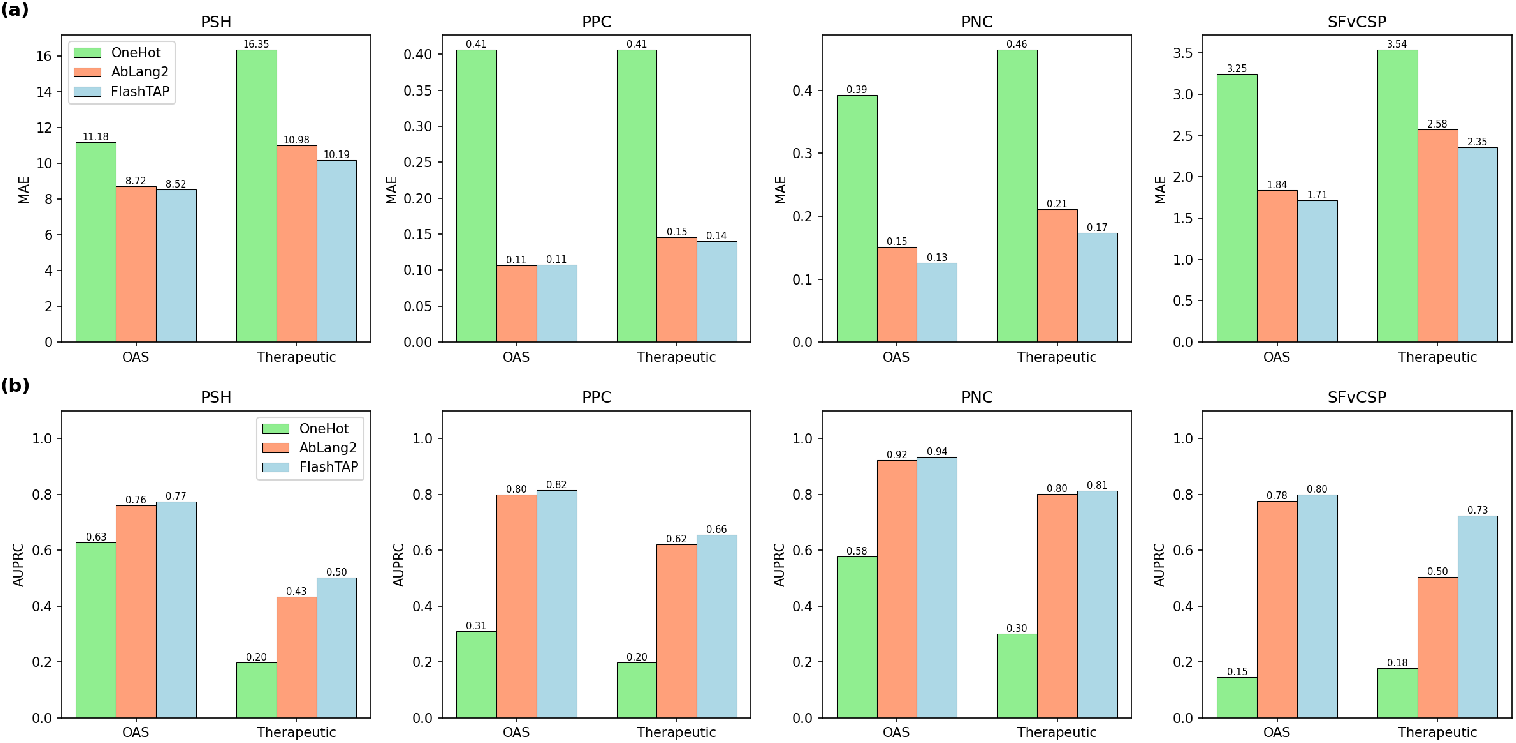
Mean average error (MAE) (a) and area under the precision recall curve (AUPRC) (b) for TAP property prediction methods. Performance for the OAS test set and therapeutic sets are shown. OneHot uses a trivial amino acid encoding. While AbLang2 provides a strong baseline, the finetuned structural FlashABB-SSS model (FlashTAP) has the best performance across all categories.

Based on the finetuned FlashABB-SSS model, we created FlashTAP, a super-fast antibody developability predictor. Flags can be assigned using the default TAP thresholds, or the probability of flags can be predicted using an isotonic regression to the OAS test set. The fit regressions are shown in Supplementary Figure F3. As shown in Figure 6, the distribution of properties between natural and therapeutic antibodies as predicted by FlashTAP qualitatively matches those generated by TAP but the agreement in flags is not perfect. Therefore FlashTAP may be used with more stringent criteria to reduce the number of false negative flag predictions. FlashTAP can predict properties for *∼*100 antibodies per second which is about 1000x as fast as TAP. This means that FlashTAP can feasibly run on the scale of repertoires or computationally generated libraries.

**Fig. 6.**
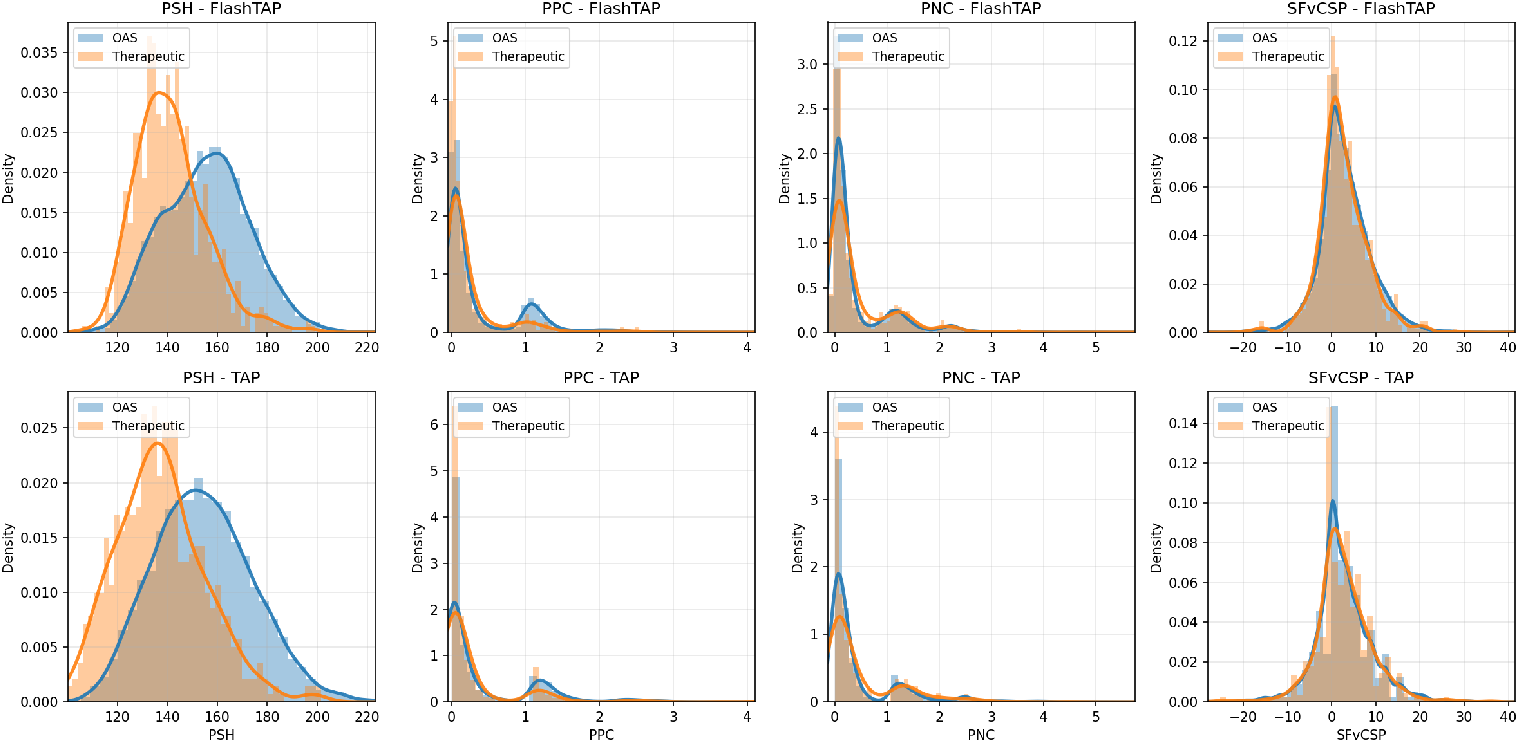
Distributional shift between natural (OAS) and therapeutic antibodies. The FlashTAP distributions qualitatively match those generated by TAP. In particular, Therapeutic antibodies have lower PSH and PPC scores.

We used FlashTAP to generate the Developability-Enriched Antibody Dataset (DevAbDab), a subset of paired OAS which is substantially less likely to have developability issues. Table 2 shows the filtering statistics using a threshold of 5% or 50% probabability of any flag.

**Table 2.**
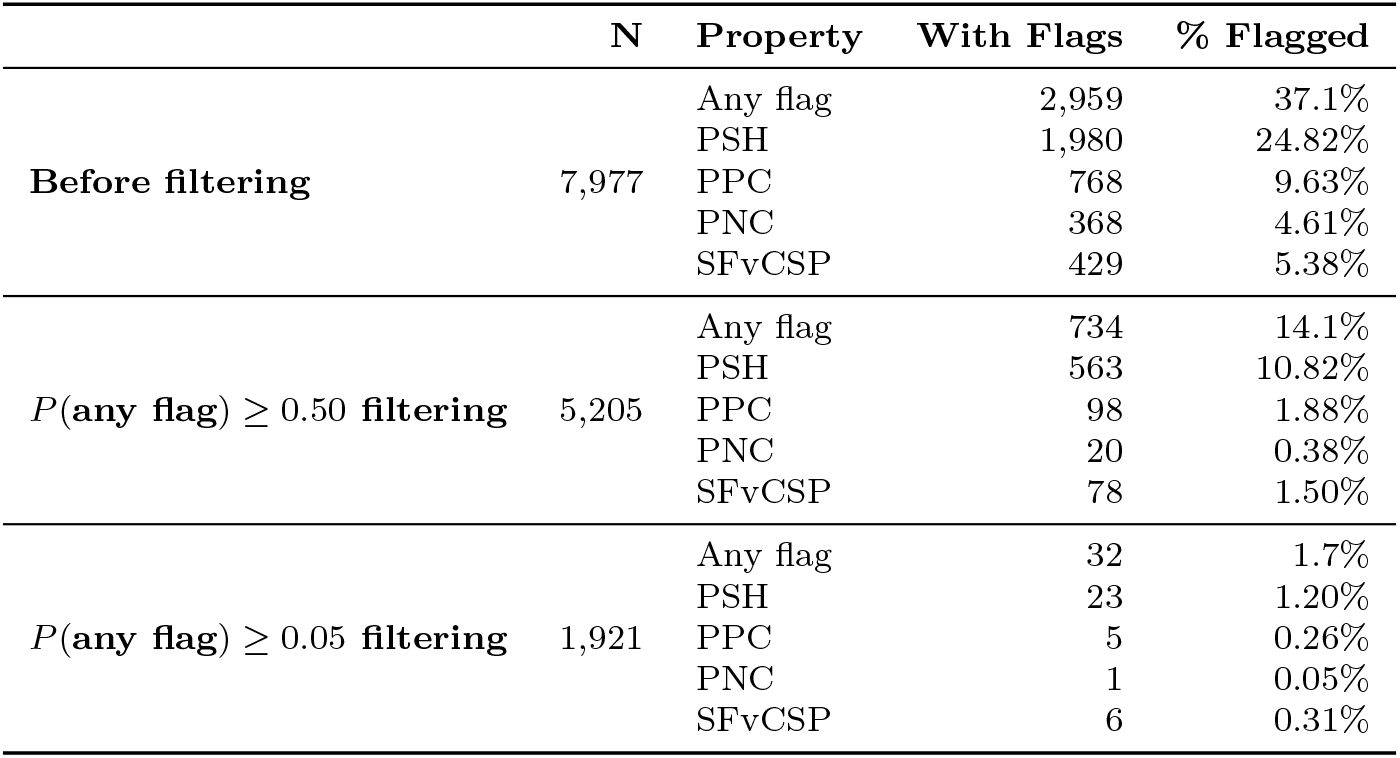
Filtering statistics on the OAS test set for FlashTAP with different filtering thresholds.

|  | N | Property | With Flags | % Flagged |
| --- | --- | --- | --- | --- |
| **Before filtering** | 7,977 | Any flag | 2,959 | 37.1% |
|  |  | PSH | 1,980 | 24.82% |
|  |  | PPC | 768 | 9.63% |
|  |  | PNC | 368 | 4.61% |
|  |  | SFvCSP | 429 | 5.38% |
| $P(\text{any flag}) \geq 0.50$ filtering | 5,205 | Any flag | 734 | 14.1% |
|  |  | PSH | 563 | 10.82% |
|  |  | PPC | 98 | 1.88% |
|  |  | PNC | 20 | 0.38% |
|  |  | SFvCSP | 78 | 1.50% |
| $P(\text{any flag}) \geq 0.05$ filtering | 1,921 | Any flag | 32 | 1.7% |
|  |  | PSH | 23 | 1.20% |
|  |  | PPC | 5 | 0.26% |
|  |  | PNC | 1 | 0.05% |
|  |  | SFvCSP | 6 | 0.31% |

Filtering at 5% reduced the number of total antibodies by about 4x but reduced the likelihood of randomly selecting a problematic antibody from 37.1% to 1.7%.

Additionally, FlashTAP predictions are differentiable, which means a flag loss gradient can be computed directly. This may enable more efficient training of models like developable pIgGen, without the need for reinforcement learning [30].

## 3 Discussion

In this work we introduce FlashABB, an antibody structure prediction method based on our new efficient IPA analog, Flashpoint Attention. FlashABB is *∼*1000x as fast as refined ABB2 predictions, *∼*7x as fast as unrefined ABB3 predictions, *∼*3x as fast as loading PDB structures with BioPython, and *∼*2x as fast as AbLang2. To our knowledge, FlashABB is the first model to accurately predict antibody structures *faster* than antibody language models can generate embeddings. We show that FlashABB predictions can be used to enrich for stability at scale and that FlashABB enables fast predictions of structural antibody features which can be leveraged for improved screening of large repertoires.

In order to implement FPA, we reformulated all internal pairwise operations as inner products, which allows us to leverage efficient attention implementations. The main benefit in this work is the improved speed, however it also uses a substantially lower memory footprint. Antibodies are relatively small, but a similar approach could be used to model other classes of proteins, in cases where the pair representation can be dropped. For example, FPA should in theory be able to accommodate extremely long context windows such as those found in natural language which could allow for structure prediction of protein complexes of tens of thousands of residues such as complete viral capsids.

The speed and memory bottleneck for FlashABB during training was the computation of the Frame Aligned Point Error (FAPE) loss. Training protein structure models for larger complexes will require the implementation of more efficient loss functions. One such solution may be to downsample the number of aligned residues in FAPE which could reduce the memory requirement from quadratic to linear in sequence length.

Together, this work establishes a new baseline in protein structure prediction: accurate prediction of meaningful structures faster than language models. While the pair representation is still necessary for general protein/complex prediction, our results show that it can be completely removed for antibodies. The efficiency of FlashABB unlocks antibody structure prediction in applications where it was previously unviable. These include repertoire modelling, real-time visualization, and training/optimization loss functions. The central innovation of FlashABB, FPA, can be adapted for future work on efficient protein structure learning. Applications of this could include inverse folding, molecular dynamics simulations, or long-context structure prediction.

## 4 Methods

### 4.1 Structure prediction: from minutes to milliseconds

As described in the introduction, there are three components of structure prediction which make it substantially slower than sequence modelling. In order of magnitude, these are coevolutionary/pair processing, molecular dynamics refinement, and inefficient kernels. Here, we outline our strategy for removing each of these barriers.

Traditional protein structure prediction relies on the analysis of coevolutionary information derived from multiple sequence alignments (MSAs) to predict putative contacts [10, 18, 21]. This information is then typically fed through a series of layers to create a pairwise representation between each pair of residues, which guides the final structure module [10]. As demonstrated in ABB2, antibody structure prediction does not need MSA processing or a sophisticated pair representation [6]. The reasons for this are twofold. First, the structural diversity of antibodies is largely limited to the complementarity determining regions, and so the contact matrix is nearly always the same. Second, antibodies “evolve” within each person to each antigen individually, so the meaningful diversity of structure is rarely represented in homologous sequences from public databases [6]. Owing to these two facts, ABB(2/3) achieve state of the art antibody structure prediction accuracy using only a trivially computed pair representation, which allows these methods to completely remove the most expensive part of structure prediction. In this work, we inherit this gain and take it a step further by removing the pair representation altogether.

The majority of time spent in ABB(2/3) is in the molecular dynamics (MD) refinement step [6]. MD refinement is particularly important for removing stereochemical errors and creating more physically plausible structures, but has limited impact on the actual backbone RMSD of the predictions [24]. In fact, in some instances, MD-refined structures may be less accurate if they are modelled in the absence of important neighbouring molecules such as waters and/or antigen chains [24]. While some applications require highly physical structures to compute physics-derived properties such as the Therapeutic Antibody Profiler [31], others only need the right backbone conformations or even benefit from a knowledge of structural contacts [2, 32]. Also, because MD refinement is typically not differentiable, any loss terms computed on MD-refined structures cannot be used to back-propagate gradients through the model. In this work, we completely remove the MD refinement step. FlashABB predictions can still be refined as needed, but we explore the potential of unrefined structures as inputs in PLM-scale applications.

### 4.2 Flashpoint attention: a novel memory-efficient analog of IPA

Methods such as ABB(2/3) demonstrate that some protein structure models do not need a sophisticated pair representation. ABB2 uses the AF2 structure module, but replaces the Evoformer with a static pair representation which contains a one-hot encoding of the chain (heavy/light) and a one-hot encoding of the relative distance between amino acid pairs, clamped between -64 and 64. ABB3 additionally includes the C*α*-C*α* distance between all pairs of residues. Even though the ABB(2/3) pair representation is trivial to compute, it still needs to be held in memory during the attention computation which requires quadratic memory and prevents the use of fast attention kernels.

Our goal is to approximate IPA under the ABB(2/3) pair representation using standard dot-product attention so that we can use fast, linear-memory attention implementations. This involves three major changes: we express the squared distances as inner products, we introduce Rotary Positional Encoding [33], and we approximate the pair values using a modular representation of position. Our Flashpoint attention (FPA) algorithm is shown in Algorithm 1 with differences to IPA highlighted in yellow. For reference, we also include a reproduction of the IPA algorithm in Appendix C.

#### Algorithm 1 Flashpoint Attention (FPA)

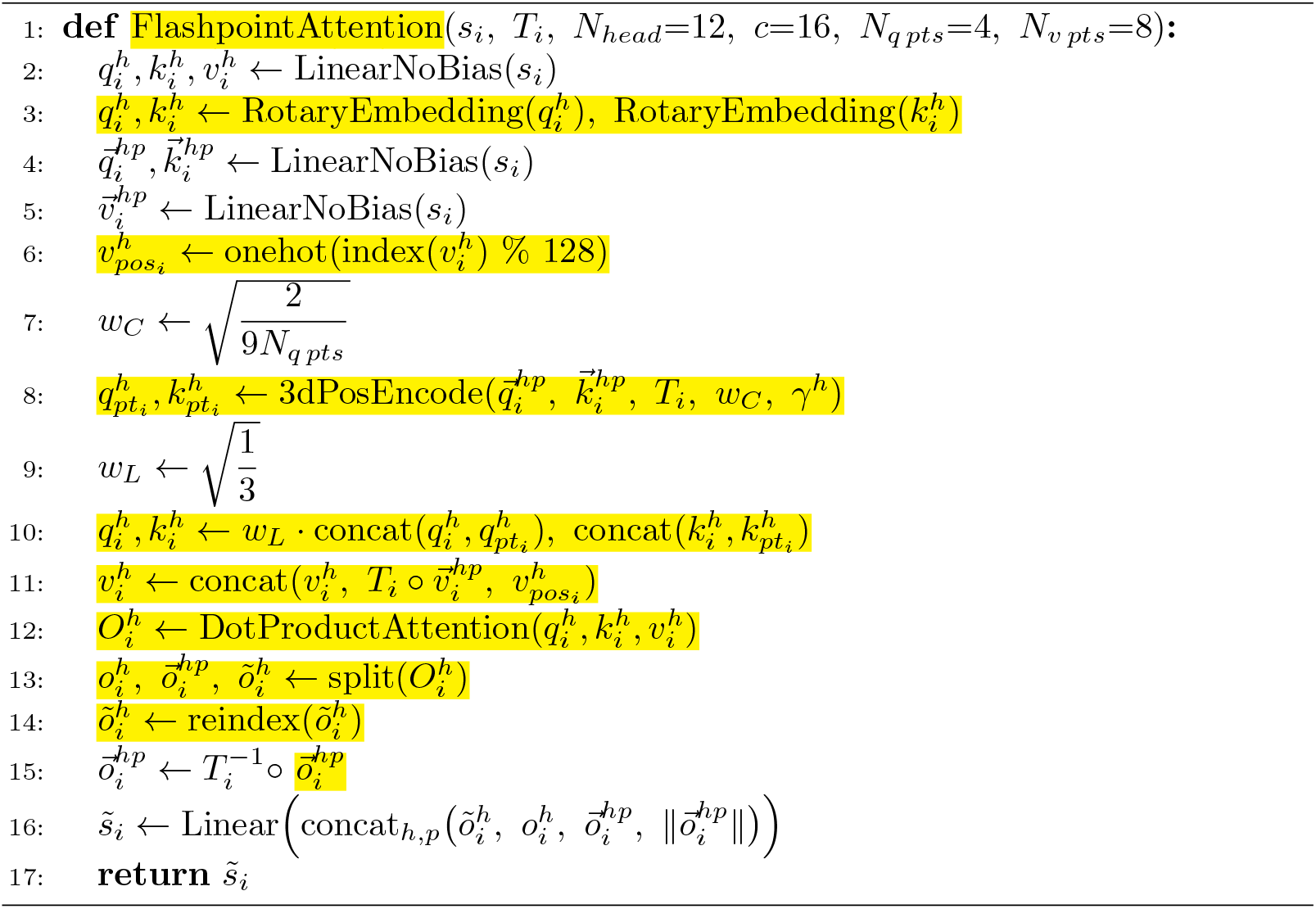

Standard IPA contains a term with the negative sum of the squared distances between virtual atoms. This serves to focus the attention on tokens with nearby atoms.

#### Algorithm 2 3D positional encoding

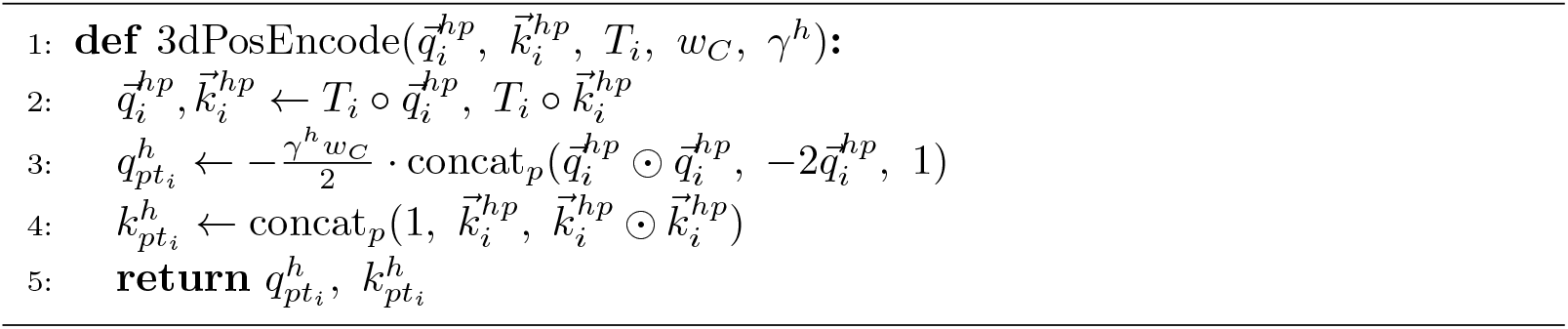

In FPA we replace this with an equivalent encoding that allows us to compute the term exactly using inner products. See Appendix D for a proof of equivalence. The advantage of computing distances using inner products is that they can be implemented using matrix multiplication which, for float32s, is about 8x as fast as non-matrix computations on modern GPUs [15].

The positional encoding in all AlphaFold-style models is handled by the bias term projected from the pair representation. In ABodyBuilder(2/3), the pair representation contains a one-hot encoding of the relative position between tokens, clamped between -64 and 64. This one-hot encoding can be linearly projected to add a positional bias of arbitrary relative positions, similar to a learned Alibi bias [34]. We experimented with replicating this exactly using FlexAttention [35], but found that Rotary Positional Encoding (RoPE) on the queries and keys converged to similar results and was faster, especially in the backwards pass [33].

IPA includes an attention-weighted sum of the pair representations in the returned value. Our implementation lacks a pair representation so we must consider what is gained in this term. The pair representation contains three terms:

The first is a one-hot encoding of whether the two chains are the same. The sum of this feature is identical to a sum of the one-hot encoded chain index, which is already included in the single representation.

The second term is the relative distance between the C*α* atoms. The sum of this is the attention-weighted average distance to the key tokens. We cannot replicate this exactly, but get an approximate measure since we get the attention-weighted average position of key tokens and then subtract the position of the query token which gives the average relative position vector.

The final term is a one hot encoding of the relative sequence position, clamped between -64 and 64. The attention-weighted sum of this term gives us a vector of the sum of the attention paid to each relative position. This can be thought of as a spectra of relative positions and may be useful for identifying, for instance, secondary structure elements. Computing this term is problematic since it contains a unique entry for each relative position, and we only produce a single value vector for each token which does not depend on the query token. To overcome this, we could append a one-hot encoding of the sequence position to the value vector for each token. This would give us a spectra of absolute positions which we could reindex to get a spectra of relative positions. However, including a one-hot vector of sequence position would necessarily have the same length as the sequence which would make our value matrix have a quadratic memory footprint in the sequence length. Instead, we append a onehot encoding of the sequence position modulus some value (in our case 128). The strategy is similar to encoding a distribution over dates by encoding the distribution over days of the week, as shown in Figure 7.

**Fig. 7.**
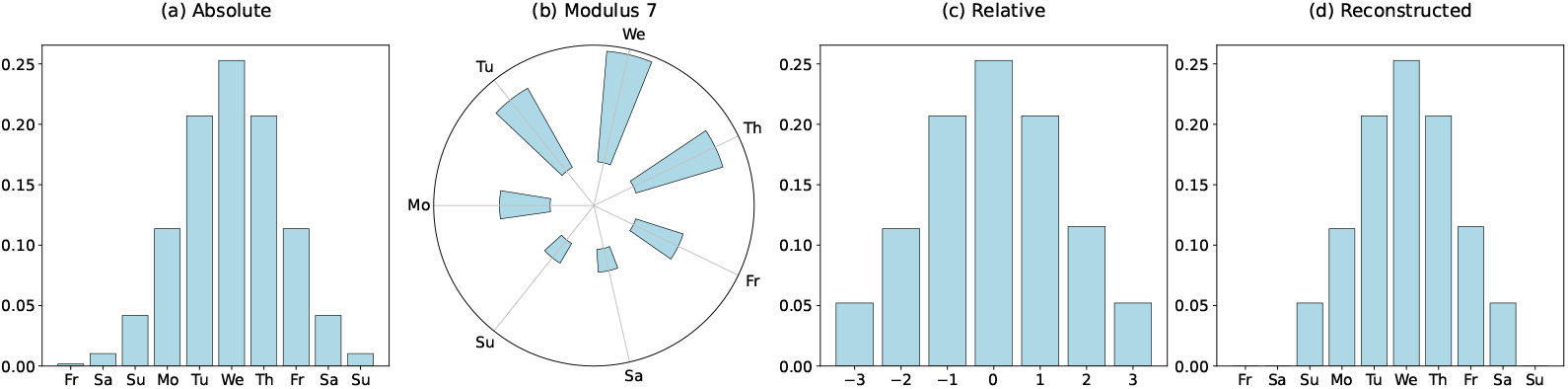
Overview of rolling position spectra demonstrated using days of the week. (a) Absolute date encoding (b) Date modulus 7 (day of the week) enables a compressed representation (c) The mod 7 representation can be shifted based on a query date to give a relative date (d) The reconstructed absolute date is reasonably accurate as long as most of the attention is paid to nearby dates.

The modular position encoding ensures that the memory remains linear, and allows us to compute a spectra of the relative positions mod 128. As long as the majority of the attention is paid to tokens which are within 64 tokens, this is equivalent to the full-sequence version. This assumption is violated in the case where tokens are attending to long-range contacts such as inter-chain contacts, but we found that the approximation still works well in practice.

### 4.3 The need for float32 precision

Squared distances between points are typically calculated like (*x*_1_ − *x*_2_)^2^. In order to factor the distance computation, we compute it using the expanded form like 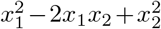 (see Algorithm 2 and Appendix D for more details). Theoretically, this is identical, but practically can suffer from numerical inaccuracies due to catastrophic cancellation [36]. We tested the extent of the inaccuracy of this method on random point clouds with coordinates drawn from [-30Å, 30Å]. For each method we compute the average error in computing the distance from a point to itself. The results are shown in Figure 2. The average error for float16s is 0.75Å whereas the average error for float32s is less than 0.01Å. An error of almost 1Å is significant since it could lead to a model attending to the wrong virtual atom. The average error of bfloat16s was the worst, at over 2Å, likely because of the reduced precision in favour of a greater range of representable numbers. Because of this, we ran all experiments using float32s. In particular, we used the default PyTorch linear memory scaled dot product attention implementation instead of FlashAttention, since FlashAttention only supports float16s.

### 4.4 Antibody structure prediction

We trained our model on the same data as ABB3, which comprises a train set of 8395 structures and a test set of 100 structures from SAbDab [24, 37, 38]. The train and test sets were split by sequence clustering and CDR similarity. We broadly followed the same training setup as ABB3, except we used a weight decay of 10^−2^, a constant learning rate of 4 · 10^−4^, and float32 precision. Removing the pair representation added some training instability which was resolved by increasing the weight decay and removing the warm restarts.

The architecture of FlashABB is identical to ABB3 except for the use of Flashpoint Attention and all usage of pair representations. In particular, FlashABB has a hidden dimension of 128, 8 layers, each with 12 heads, and a head dimension of 16. Both FlashABB and ABB3 have approximately 7.2M parameters.

### 4.5 Fast stability prediction

Fast structure prediction unlocks fast prediction of properties which depend on structure. To demonstrate this, we predicted the ProteinMPNN likelihood of antibodies conditioned on their true sequence and FlashABB-predicted structure (see Figure 8). As has been shown for general proteins, we found that these likelihoods are correlated with experimentally measured melting temperatures. We can run this method on *∼*50 antibodies per second which enables stability enrichment for libraries of millions of antibody sequences. More details can be found in Appendix E.

**Fig. 8.**
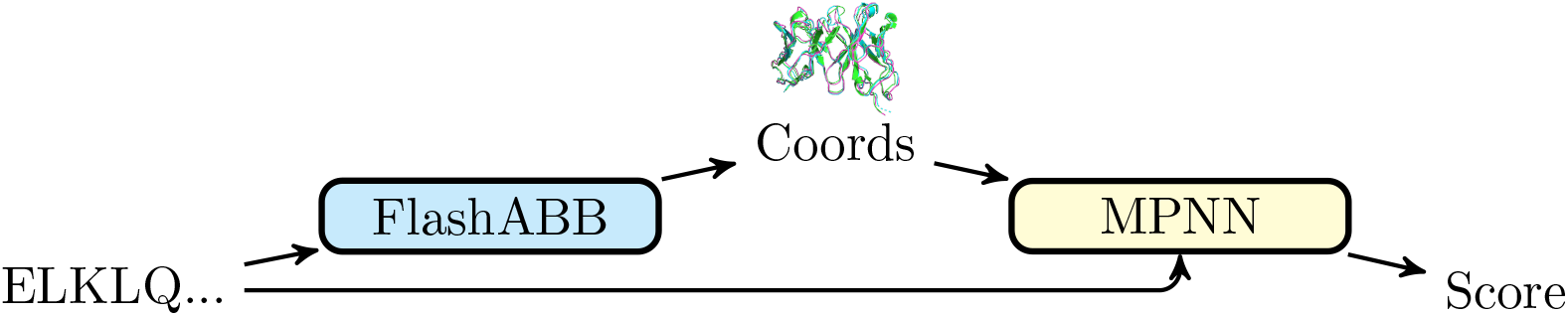
Structure as a latent representation for a sequence model. Antibody structures are predicted and the backbone coordinates are fed into an inverse folding model to predict the likelihood of each residue.

### 4.6 Structure as a latent representation for language modelling

As FlashABB is faster than antibody language models, there is little practical overhead to using structure as an input for antibody language modelling. We demonstrated this by creating a seq2seq model for antibodies which runs through modelled structure as a latent representation. The resulting model, which we call seq2struct2seq maintains the benefits of inverse folding models without the computational overhead. The model uses FlashABB to generate a backbone structure which is passed, alongside partially masked tokens, to FlashABB-inv. The architecture of FlashABB-inv is identical to FlashABB except that it does not perform backbone structure updates and ends with a sequence logit prediction head. We call the combined model (FlashABB -¿ FlashABB-inv) FlashABB-SSS (seq2struct2seq). During training, we freeze the weights of FlashABB. We randomly split the entirety of paired OAS (*∼*3M sequences) into training and validation sets, and trained FlashABB-SSS to perform masked token prediction using a standard setup [39, 40]. In particular, we masked 15% of tokens which were randomly swapped to [MASK] (80%), self (10%), or a random amino acid (10%).

FlashABB-SSS achieves lower perplexity and higher sequence recovery than regular antibody language models. However, this can be largely attributed to its ability to extract amino acid identities from subtleties in the backbone coordinates, which constitutes a form of data leakage. Despite this, we found that FlashABB-SSS, was a useful pretrained model to finetune for property prediction. To demonstrate this, we trained FlashTAP, a model which predicts the physiochemical properties of antibodies used for developability prediction by TAP [7]. We trained FlashTAP by finetuning FlashABB-SSS, followed by an MLP and sum layer to predict Patches of Surface Hydrophobicity, Positive Charge, and Negative Charge (PSH, PPC, PNC) as well as the Structural Fv Charge Symmetry Parameter (SFvCSP). We also trained an equivalent model by finetuning AbLang2, an antibody language model trained on OAS. Both models were trained on the set of *∼*80K precomputed ABB2-predicted natural antibody structures released alongside ImmuneBuilder, labeled by TAP2 and randomly split into 80%/10%/10% train/test/val splits [6, 7]. The train/val/test splits were random but we observed a low level of CDR similarity between the train and test sets and only a modest performance difference on test sequences with high similarity to train. More details are available in Appendix F. We evaluate performance on the test set of natural antibodies from OAS as well as the entire therapeutic set.

## Appendix A Comparison to Liu et al. [19]

Concurrently with our work, Liu et al. [19] developed Flash IPA, another method for implementing IPA using inner products which enables fast, memory efficient IPA. As in our work, FlashIPA augments key and query vectors to compute the sum of the squared distances of the virtual atoms and augment the value vectors to compute the attention-weighted sum of the relative positions.

In contrast to Liu et al. [19], we do not factor the pair representation. The reason for this is twofold. First, it is unclear in general that pair representations can be effectively factored into low-rank matrices. If this were possible, pair representations could always be learned using standard keys and queries of sufficient size. Second, factorizing an *n* × *n* pair representation requires realizing that representation in memory, which is inherently quadratic. To overcome this, Liu et al. [19] only compute the pair representation for the k-nearest neighbours which, for structure prediction/generation models, can only be done in sequence space.

Instead, we introduce rolling position spectra to overcome the key pair representation contribution for antibody modelling. This is a good approximation of the relative position values for attention paid to local residues in sequence space, for arbitrarily long sequences. However, our approach is limited to trivial pair representations which may be limiting for other applications. We leave an investigation into similar efficient approximations of more sophisticated pair representation attributes for future work.

Additionally, we do not actually use FlashAttention. This is because FlashAttention only supports float16s and our experiments indicate that these suffer from catastrophic cancellation under the new distance calculation (see Figure 2). Instead, we use the default memory efficient scaled dot product attention in PyTorch which still results in greatly improved speed and linear memory.

## Appendix B Test antibody sequences

For reference, the sequences used to test FPA properties are listed here

Heavy:

~~~
QVQLVQSGAEVKKPGSSVKVSCKASGGTFSSLAISWVRQAPGQGLEWMGGIIPIFGTANY
AQKFQGRVTITADESTSTAYMELSSLRSEDTAVYYCARGGSVSGTLVDFDIWGQGTMVTV
SS
~~~

Light:

~~~
DIQMTQSPSTLSASVGDRVTITCRASQSISSWLAWYQQKPGKAPKLLIYKASSLESGVPS
RFSGSGSGTEFTLTISSLQPDDFATYYCQQYNIYPITFGGGTKVEIK
~~~

## Appendix C Invariant Point Attention

For reference, here we reproduce the Invariant Point Atteniton algorithm from Jumper et al. [10].

### Algorithm 3 Invariant Point Attention (IPA)

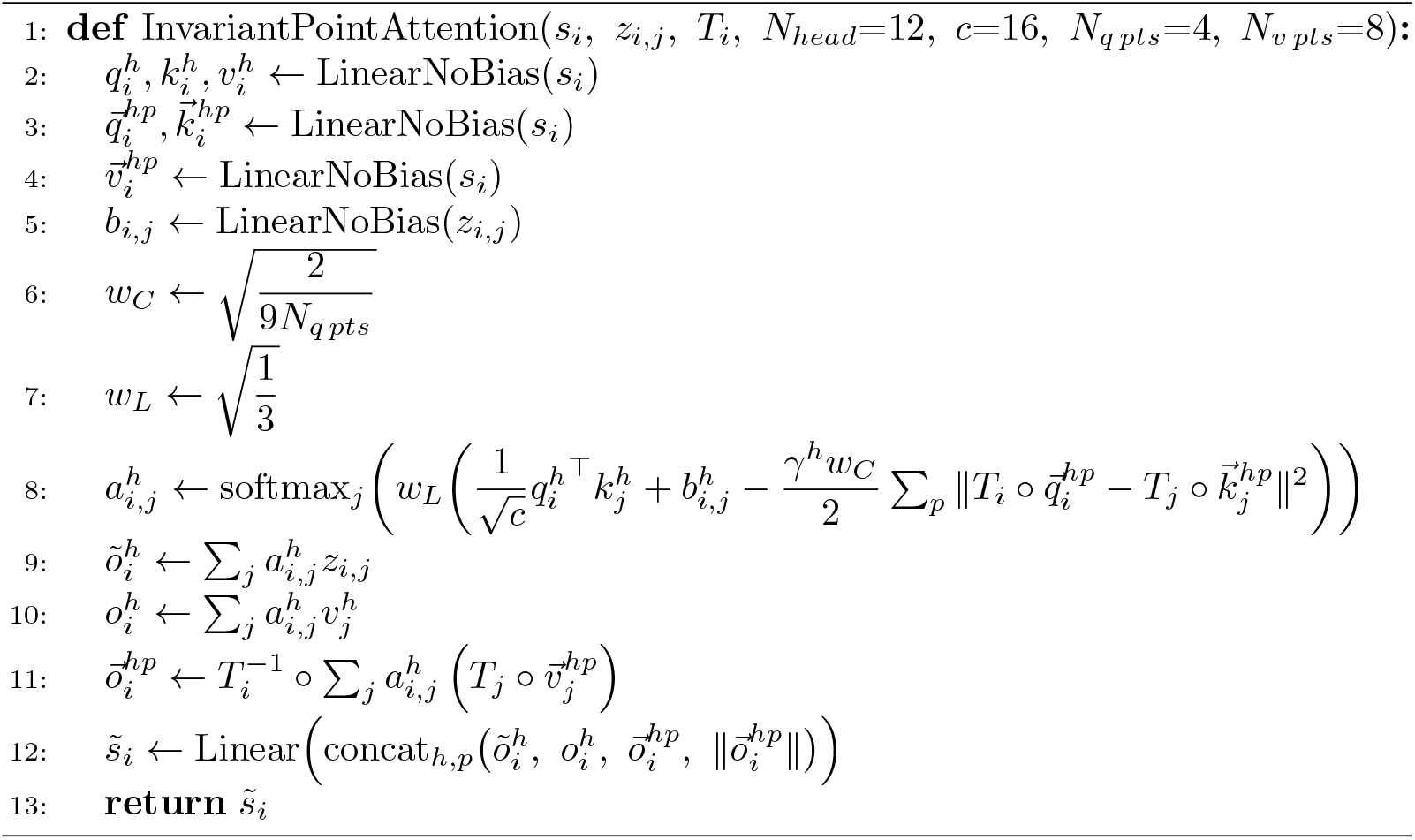

## Appendix D Proof of equivalence in distance computation

### Theorem 1

*Consider* 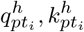 *as defined on line 6 of Algorithm 1 and* 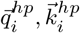 *as defined in Algorithms 3 and 1. Then:*

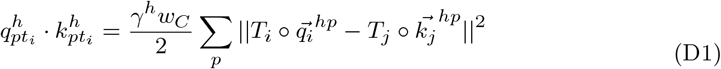

*In particular, the relative distance term of IPA can be expressed as an inner product*.

*Proof*

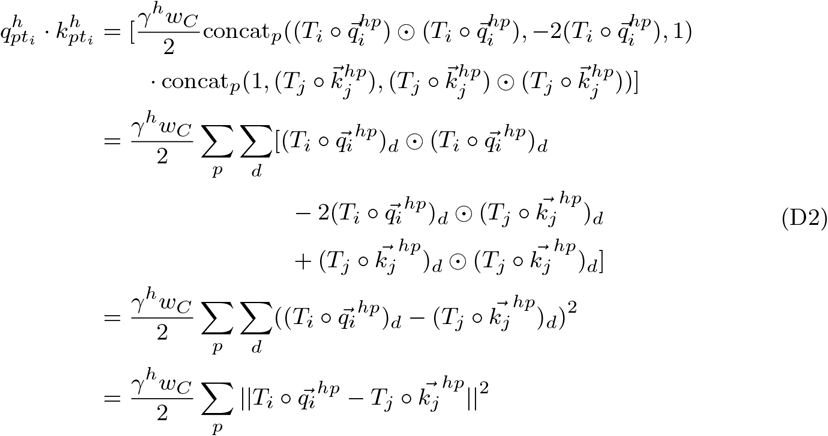

## Appendix E Stability prediction details

For antibody stability prediction we compute each structure using FlashABB and then run ProteinMPNN conditioned on the predicted structure. To estimate the relative stability, we compute the log likelihood of the ProteinMPNN logits with two modifications. First, we use the average log probability of each amino acid rather than the sum to avoid penalizing longer antibodies. Second, we clamp the maximum probability at 10%. This removes noise from the model deciding between plausible residues and focuses only on mutations which are clearly deleterious.

## Appendix F FlashTAP

To ensure that the performance of FlashTAP was not being driven by memorizing training samples, we computed the maximum concatenated CDR sequence identity to a training sequence for every sequence in test using MMSeqs2 [41]. The results are shown in Figure F1 Less than 3% of test sequences had a sequence in train with at least 90% CDR sequence identity. Additionally, the mean average error was only modestly different between bins of maximum similarity to train.

## Appendix G Inverse folding by inverting folding

Inverse folding models are designed to predict a sequence conditioned on a specified fold. However, inverse folded models are not directly optimized to predict a sequence which maximizes the likelihood of refolding into the desired fold. Additionally, inverse folding models can be sensitive to small perturbations in the reference structure. To overcome these limitations, methods like BindCraft perform gradient descent on a structure prediction model to optimize the refolding objective directly [42]. However, these methods are typically very computationally intensive because they need to compute the forward and backward pass of a structure prediction model many times. Here, we show that FlashABB enables improved inverse folding by jointly optimizing a sequence on the AbMPNN loss and the FlashABB target structure RMSD [8]. A diagram is shown in Figure G4. Because FlashABB is so fast, we can compute 50 iterations of this optimization in a few seconds per antibody.

**Fig. F1.**
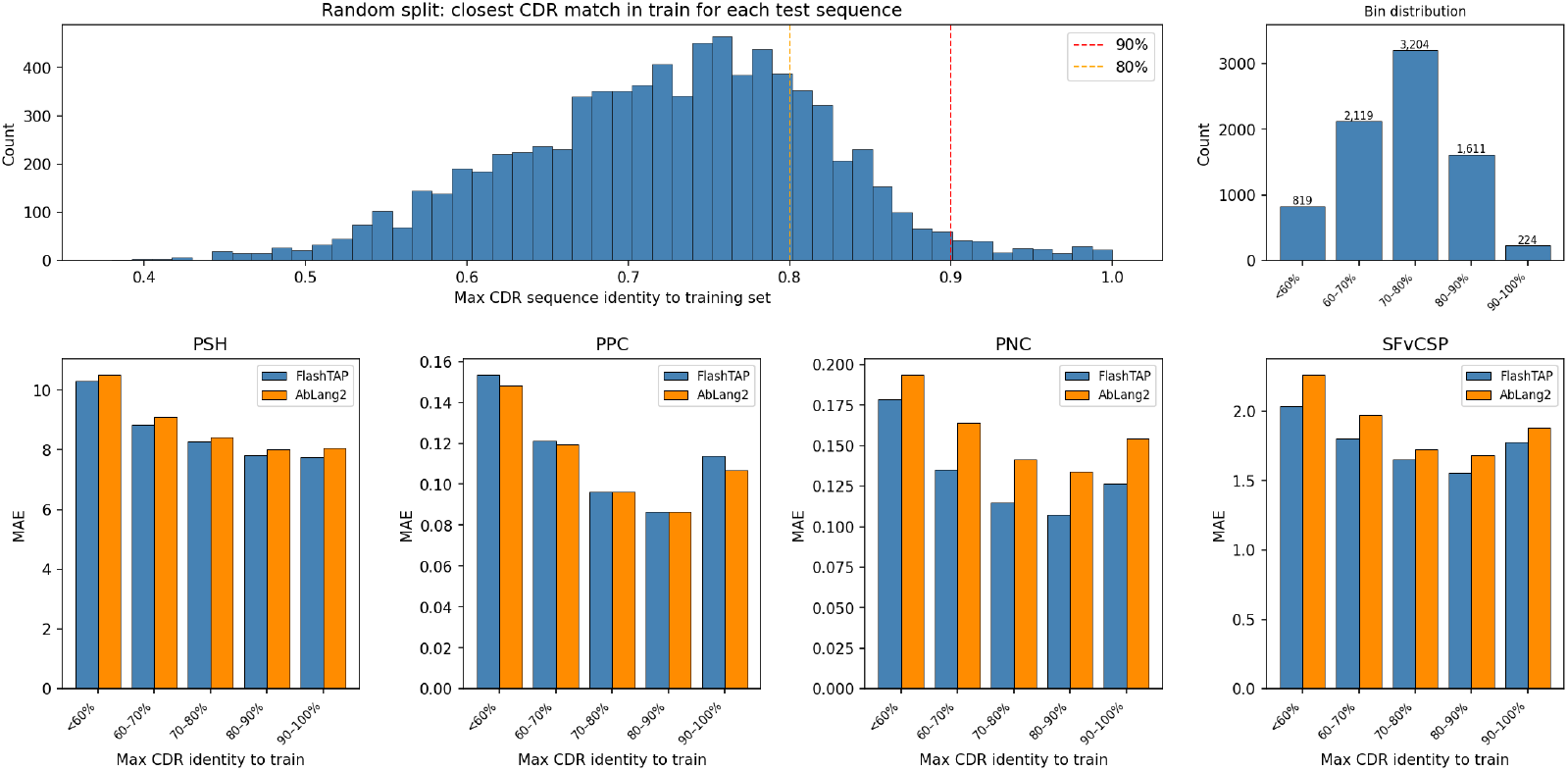
Maximum concatenated CDR similarity to train for TAP property test sequences. Less than 3% of sequences have a highly similar sequence in train and these sequences do not show a dramatically lower error in any category.

We optimize a sequence by following the gradient of the AbMPNN loss on the target structure and the RMSD of our current predicted structure to the target structure. We maintain a “soft” representation which is cast to a “hard” representation when we need token labels by taking the argmax. The sequence representation is seeded with the AbMPNN-predicted sequence. This representation is converted to a probability distribution over amino acids at a high temperature which yields a fairly uniform representation. The distribution then iteratively converges to a one-hot optimized representation by following the gradient of the combined loss and sampling at increasingly lower temperatures.

**Fig. F2.**
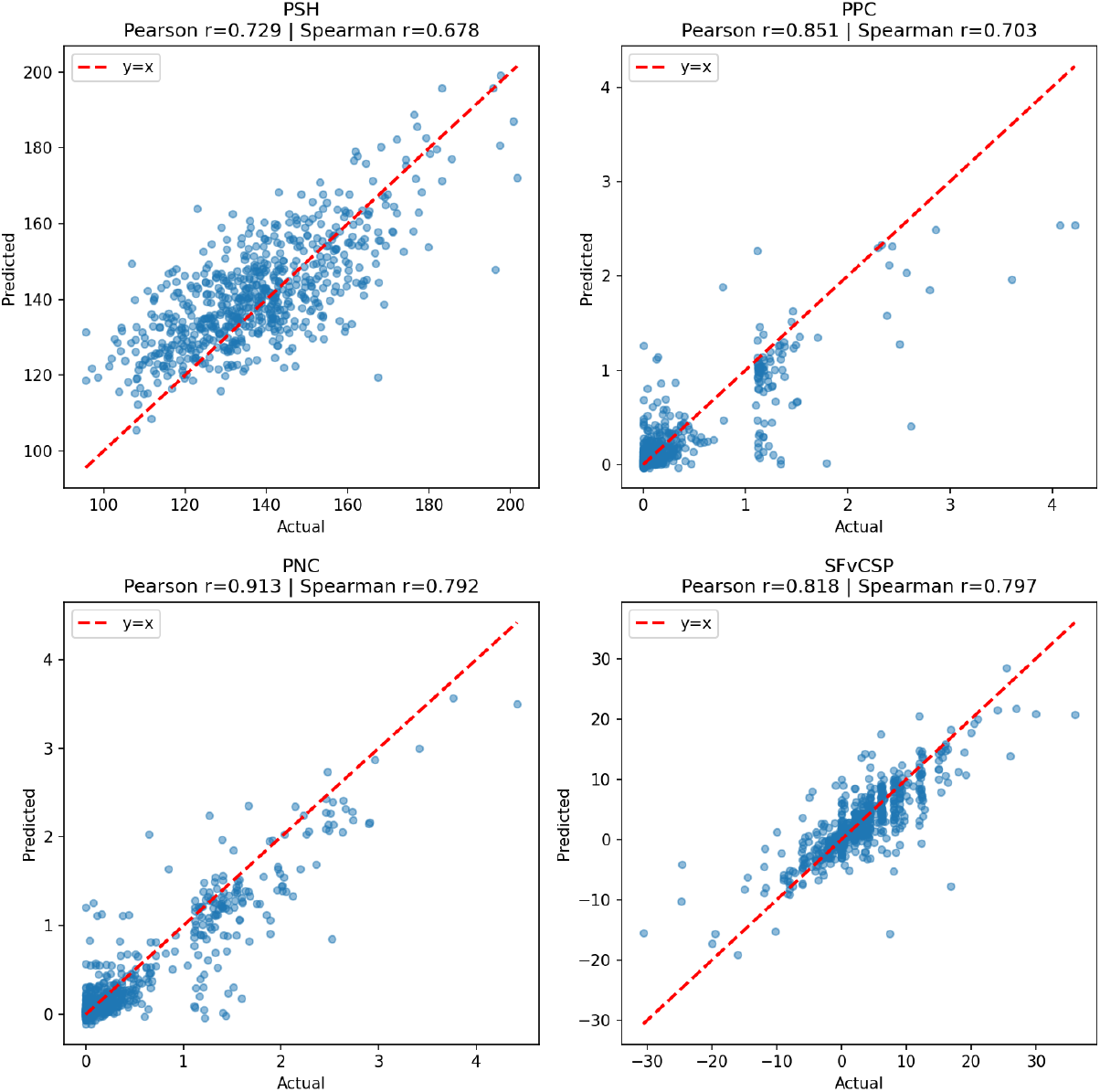
FlashTAP-predicted properties vs TAP2 computed properties evaluated on the therapeutic set. The plots qualitatively match the ABB1 vs ABB2 plots reported in Raybould et al. [7].

### Algorithm 4 Antibody Inverse Folding via Gradient Descent

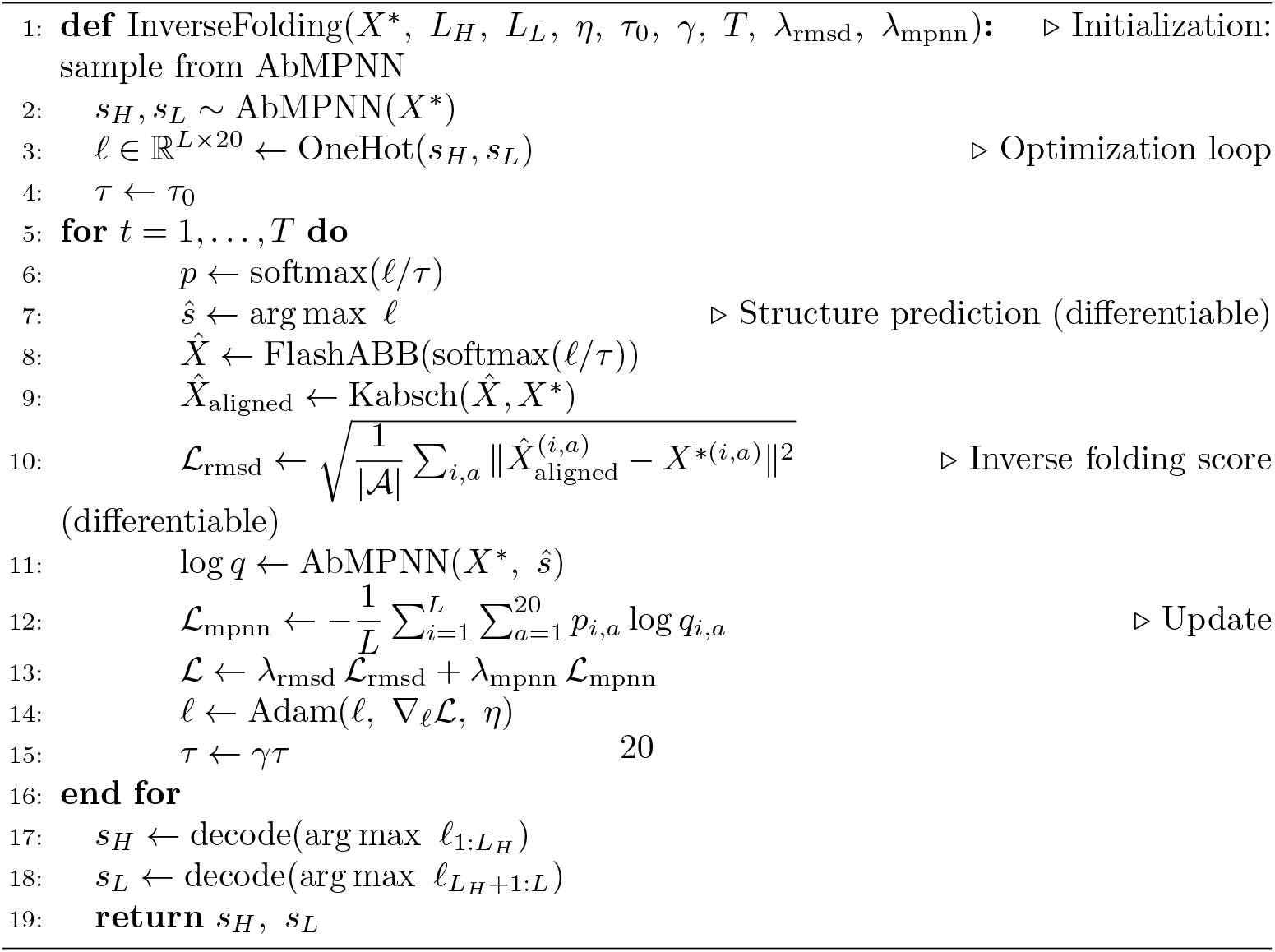

**Fig. F3.**
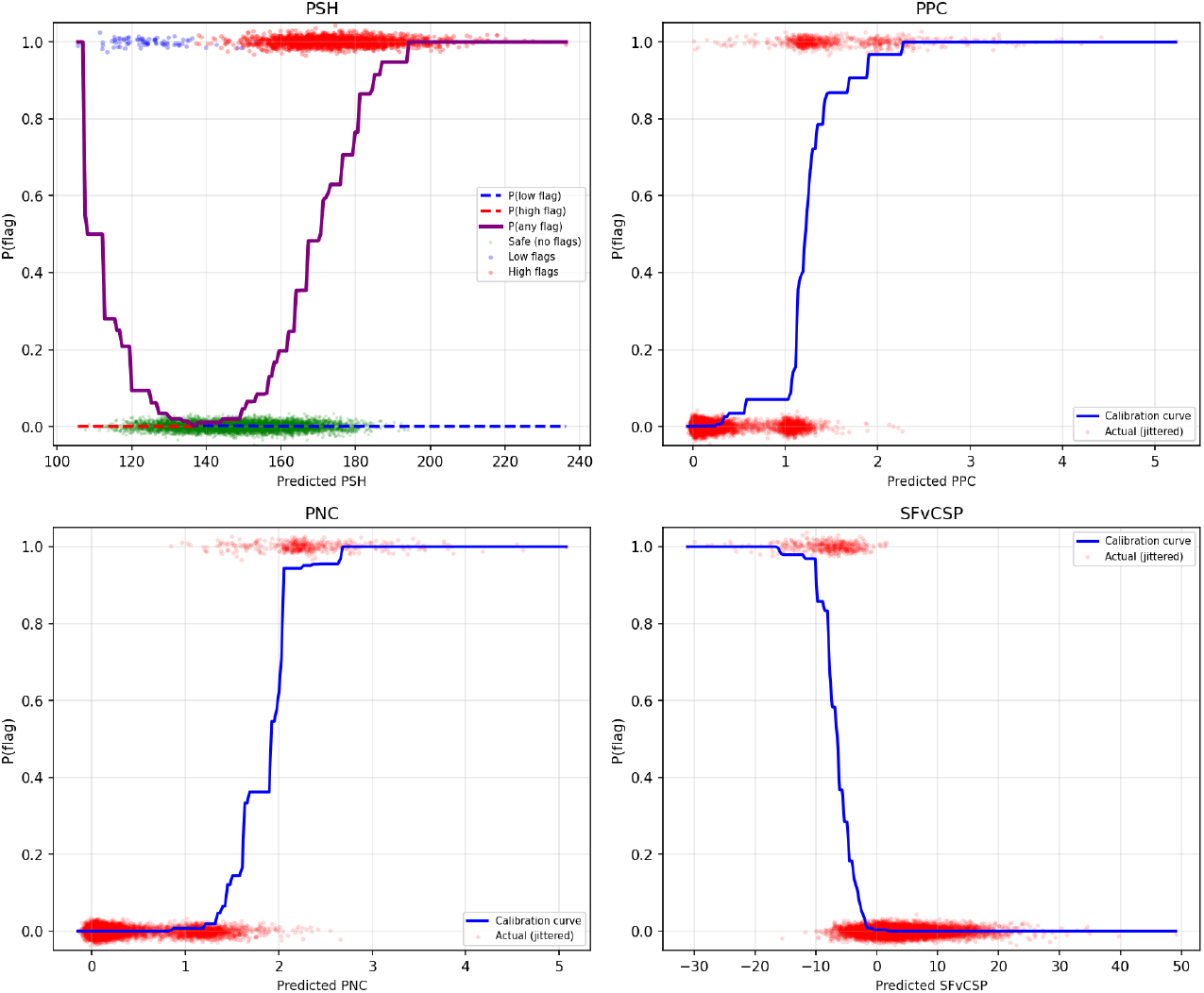
Isotonic calibration curves for mapping the FlashTAP regression to values to flag probabilities. For PSH, the high and low flags are calibrated separately and the product of 1-P(no flag) is shown.

We tested our iterative inverse folding optimization method on the test set of FlashABB to ensure that there was no data leakage of sequence into the predicted structures. We compare the AbMPNN + FlashABB-optimized sequences to the AbMPNN only sequences. The results are shown in Figure G5.

As expected, the FlashABB-predicted RMSDs to the ground-truth were substantially lower for the FlashABB-optimized sequences. In the FlashABB optimization approach we directly minimize the RMSD, so it could be that these sequences were simply exploiting biases in FlashABB which are unrelated to the physical similarity to the target. We also looked at the performance broken down by region on the antibody. The results are shown in Figure G6.

**Fig. G4.**
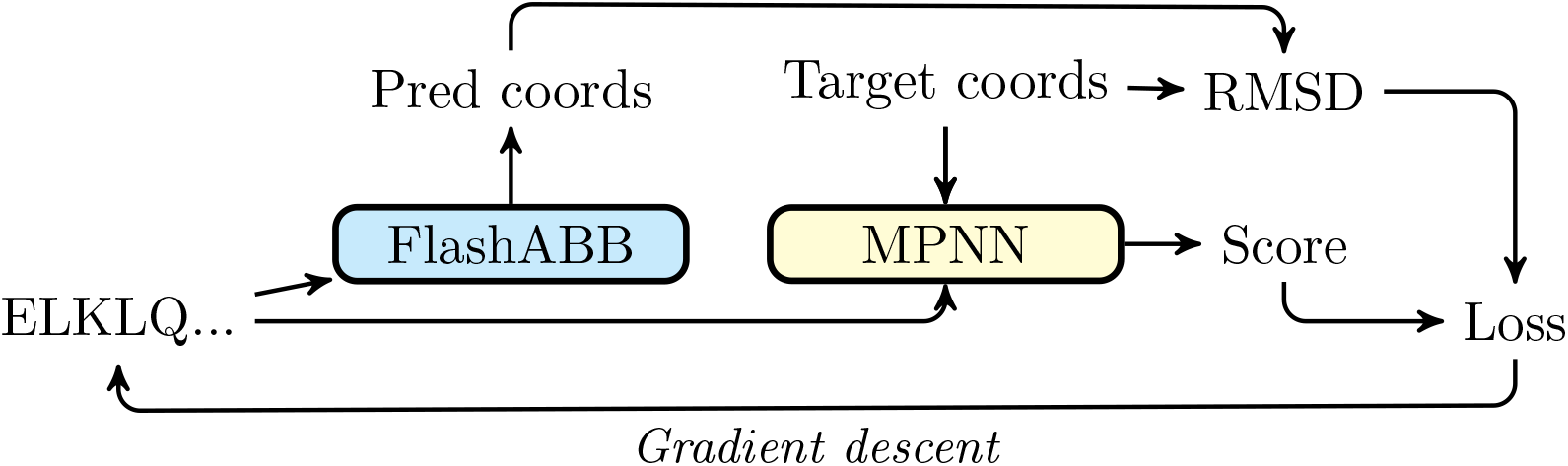
Inverse folding by gradient descent. We compute a soft representation of the sequence which is fed into FlashABB and AbMPNN (with the target coordinates). The loss is the AbMPNN log likelihood minus the RMSD between the predicted and target structures. By optimizing the sequence over the gradient of this loss we can iteratively move towards sequences which are more likely to adopt the desired structure.

**Fig. G5.**
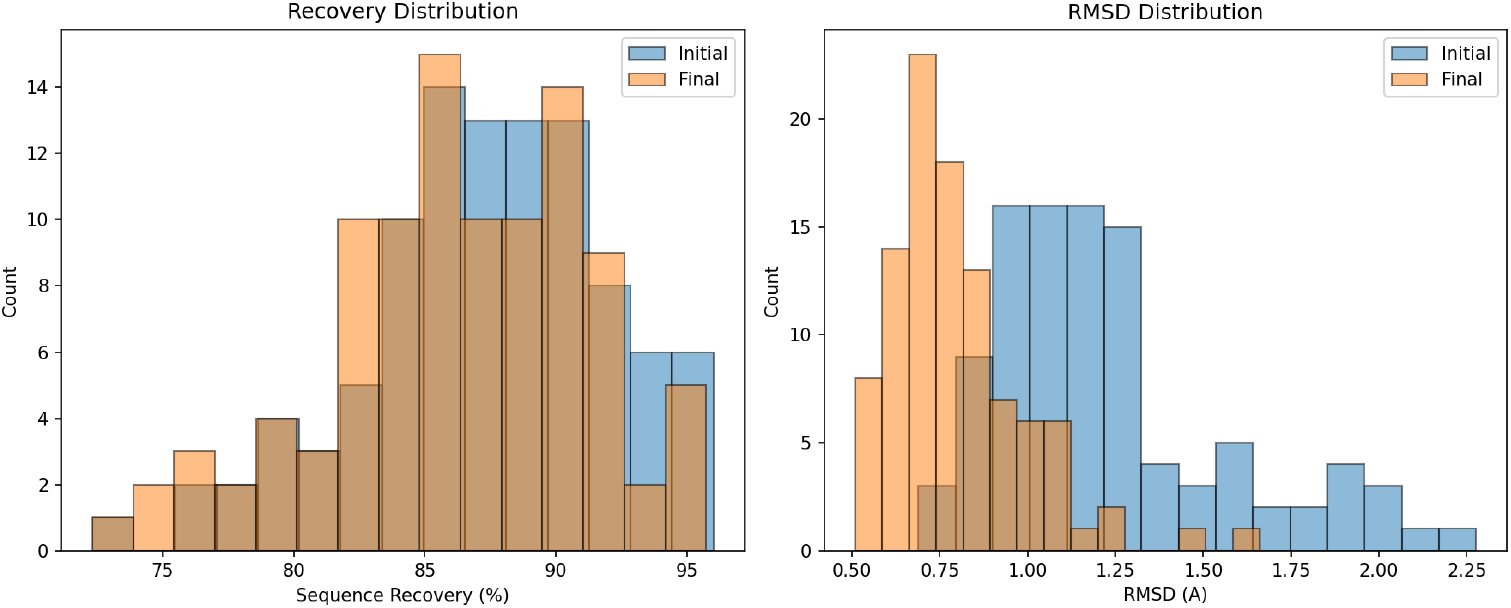
Change in sequence recovery and RMSD for FlashABB-optimized antibodies compared to the AbMPNN starting sequences. Optimization substantially improves predicted RMSD but slightly reduces sequence recovery.

**Fig. G6.**
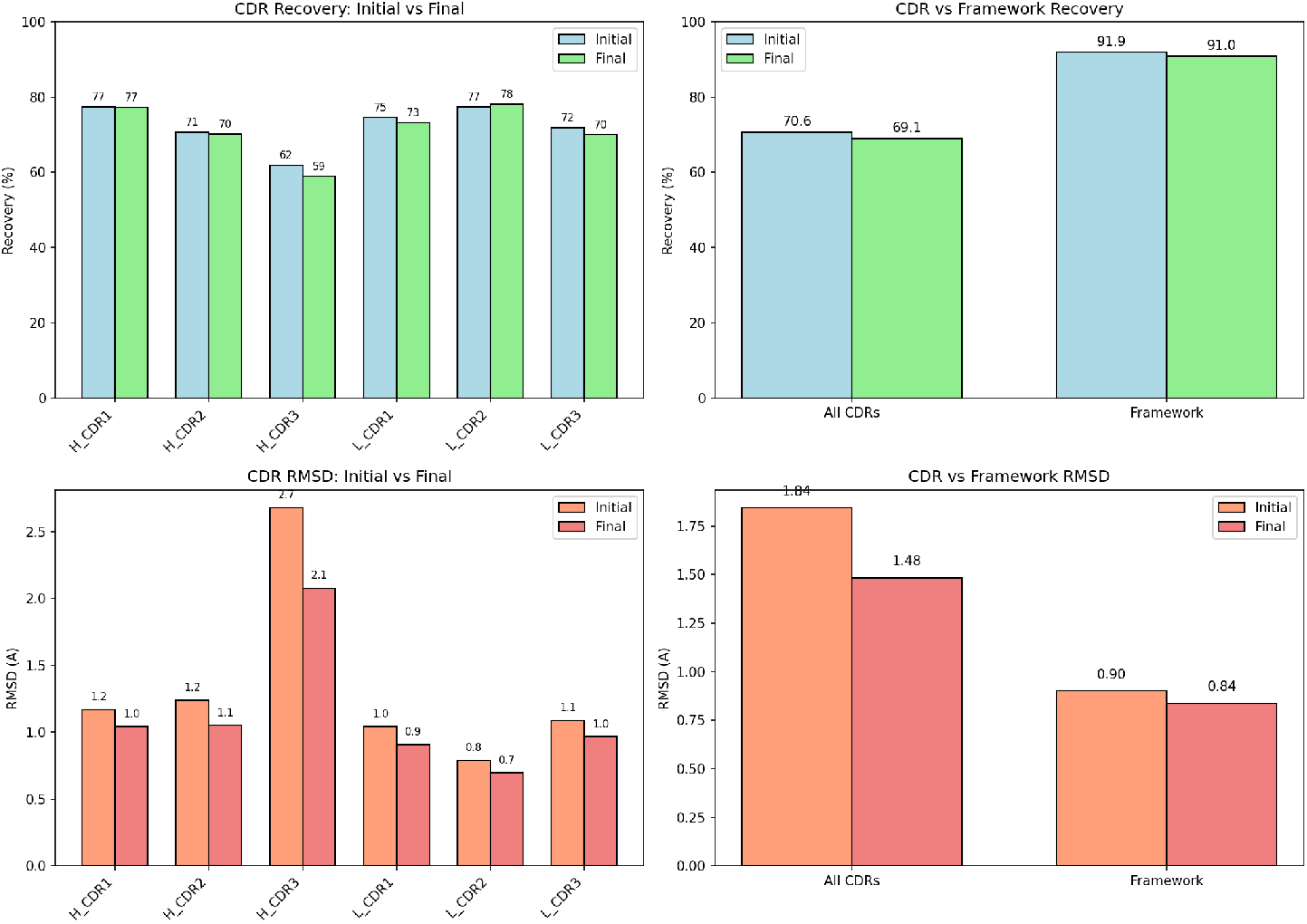
Average sequence recovery and predicted RMSD broken down by region.

## Notes

https://github.com/oxpig/FlashABB

### Competing Interest Statement

The authors have declared no competing interest.

### Funding

- Engineering and Physical Sciences Research Council: EP/S024093/1

